# Immunomodulatory properties of umbilical cord blood-derived small extracellular vesicles and their therapeutic potential for inflammatory skin disorders

**DOI:** 10.1101/2021.08.06.455399

**Authors:** Sílvia C. Rodrigues, Renato M. S. Cardoso, Patricia C. Freire, Cláudia Gomes, Filipe Duarte, Ricardo Neves, Joana Simões-Correia

## Abstract

Umbilical cord blood (UCB) has long been seen as a rich source of naïve cells with strong regenerative potential, likely mediated by small extracellular vesicles (sEV). More recently, small extracellular vesicles (sEV), such as exosomes, have been shown to play essential roles in cell-to-cell communication, via the transport of numerous molecules, including small RNAs. Often explored for their potential as biomarkers, sEV are now known to have regenerative and immunomodulating characteristics, particularly if isolated from stem cell-rich tissues. In this study, we aim to characterize the immunomodulating properties of umbilical cord blood mononuclear cell sEV (herein referred as Exo-101), and explore their therapeutic potential for inflammatory skin diseases. Exo-101 was shown to shift macrophages toward an anti-inflammatory phenotype, which in turn exert paracrine effects on fibroblasts, despite previous inflammatory stimuli. Additionally, the incubation of PBMC with Exo-101 resulted in an reduction of total CD4^+^ and CD8^+^ T-cell proliferation and cytokine release, while specifically supporting the development of regulatory T-cells (Treg), by influencing FOXP3 expression. In a 3D model of psoriatic skin, Exo-101 reduced the expression of inflammatory and psoriatic markers IL-6, IL-8, CXCL10, COX-2, S100A7 and DEFB4. *In vivo*, Exo-101 significantly prevented or reversed acanthosis in imiquimod-induced psoriasis, and tendentially increased the number of Treg in skin, without having an overall impact on disease burden. This work provides evidence for the anti-inflammatory and tolerogenic effect of Exo-101, which may be harnessed for the treatment of Th17-driven inflammatory skin diseases, such as psoriasis.

## INTRODUCTION

Virtually every living cell releases extracellular vesicles (EV), which can be classified based on size and marker expression. One of the smallest known groups of EV, exosomes, have a diameter typically ranging from 30 to 100 nm and originate from endosomes (Raposo and Stoorvogel 2013). Composed of lipids, proteins and nucleic acids, these ubiquitous vesicles are thought to be involved in multiple diseases, including inflammatory and autoimmune skin conditions (Wang et al. 2019). Due to their physical characteristics, which allow them to carry molecules across long distances, EV are often explored for their potential as biomarkers (McBride et al. 2017; Wang et al. 2020). Physiologically, small EV (sEV), such as exosomes, are key mediators of cellular communication, namely through microRNAs (van Niel et al. 2018). Hence, depending on the producing cell, sEV may have modulating characteristics, which can potentially be harnessed for therapeutic purposes. In fact, these naturally-produced vesicles are currently explored for the treatment of several conditions, including wound healing (Henriques-Antunes et al. 2019) and autoimmune diseases (Bai et al. 2017; Kim et al. 2005; Ma et al. 2019). Their use replaces cell therapies (Phinney and Pittenger 2017; Di Trapani et al. 2016), while conferring advantages, namely concerning handling and formulation.

Umbilical cord blood (UCB) is a rich source of stem cells and immature T-cells (Harris et al. 1992) with potent suppressive ability (Godfrey et al. 2005). The collection of UCB, commonly seen as medical waste, is non-invasive and has limited ethical concerns. We have previously shown that sEV from UCB mononuclear cells (MNC) accelerate the healing of diabetic wounds (Henriques-Antunes et al. 2019). In this study, we characterize the immunomodulating properties of UCB-MNC-derived sEV, Exo-101, and explore their potential in the treatment of psoriasis symptoms.

## RESULTS

### Exo-101 has anti-inflammatory and tolerogenic effects, modulating different immune players directly and indirectly

While exploring the therapeutic potential of Exo-101 for chronic wound healing (Cardoso et al. 2021; Henriques-Antunes et al. 2019), we found differences in the immunological profile of treated versus control skin. Namely, 194 genes associated with immune system processes or inflammatory responses, corresponding to about 16% of all measured genes, were differentially expressed between the two groups of animals (Figure S1). Based on these results and on existing literature, we performed a series of *in vitro* proof-of-concept experiments to determine the nature and strength of Exo-101’s immunomodulatory effect.

In line with studies using sEV from different sources (Cao et al. 2019; Hu et al. 2020), Exo-101 promotes the differentiation of macrophages into an anti-inflammatory M2, rather than a pro-inflammatory M1 phenotype (Figure 1). This effect is present both in unstimulated (Figure 1a-c) and LPS-stimulated macrophages (Figure 1d-f) and correlates with the *de novo* synthesis and release of inflammatory mediators (Figure 1a-l). Specifically, Exo-101 administration significantly decreases the level of *IFNG, IL1B, PTSG2* and *TNFA* mRNA in macrophages, as well as the release of TNFα and CCL20 proteins. These data strongly indicate that Exo-101 plays a direct role in macrophage regulation, having an anti-inflammatory effect, even in the context of an acute pro-inflammatory stimulus.

**Figure 1:**
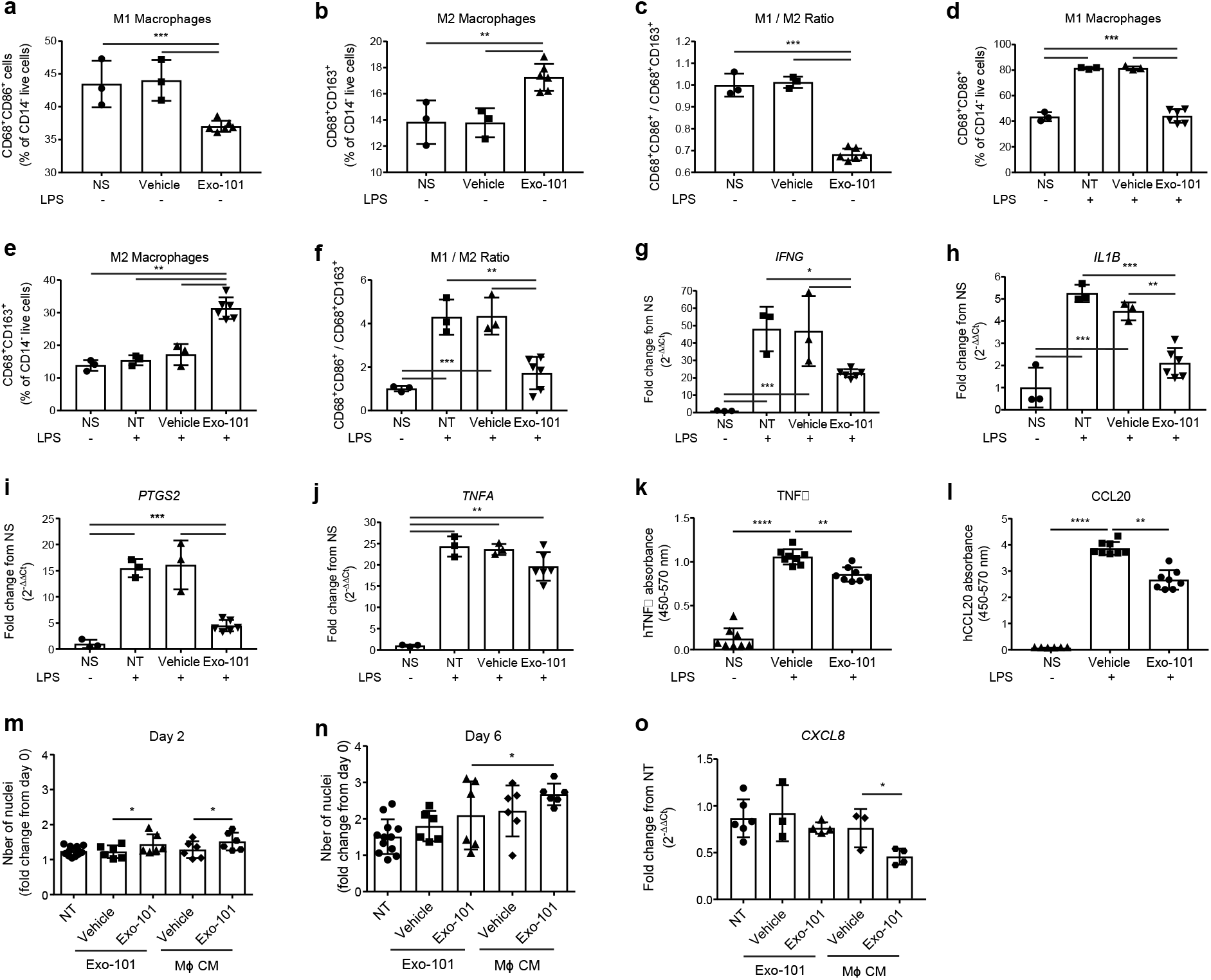
Exo-101’s modulatory effects on macrophages and fibroblasts *in vitro*. (**a-f**) THP-1-derived macrophages, with or without LPS stimulation as indicated, were incubated with 1×10^10^ particles/mL of Exo-101 for 24h, before flow cytometry analysis (n≥3). (**g-j**) Relative expression of M1-associated genes by qPCR (n≥3) and (**k, l**) release of pro-inflammatory molecules by ELISA (n≥6). (**m, n**) Proliferation of normal human dermal fibroblasts, 2 and 6 days after incubation with Exo-101 (1×10^10^ particles/mL) or media of Exo-101-stimulated macrophages (n≥6), and (**o**) CXCL8 expression at 48h (n≥3). All results are presented as mean +/- SD. *p≤0.05, **p≤0.01, ***p≤0.001, ****p≤0.0001. NS, non-stimulated; NT, non-treated; CM, conditioned media.

At the local level, Exo-101 directly promotes the proliferation of dermal fibroblasts (Figure 1m-n). Interestingly, the incubation of fibroblasts with conditioned medium from Exo-101-stimulated macrophages likewise modulates cell proliferation, as well as chemokine synthesis, with a more pronounced effect when compared to direct Exo-101 stimulation (Figure 1o). These results suggest that, while Exo-101 does act directly on target skin cells, its effects are likely reinforced by indirect mechanisms, depending on neighboring immune cell populations, such as macrophages.

To evaluate the response of T-cells to Exo-101 stimulation, in a physiological context, whole human PBMC were incubated for 6 days with a single dose of 1×10^10^ vesicles/mL, following activation with α-CD3/-CD28. Exo-101 significantly reduced the proliferation of total, CD4^+^ and CD8^+^ T-cells, as well as the intracellular content of IFNγ in each of these populations (Figure 2a-f). The release of TNFα and CCL20 by total PBMCs was also significantly decreased (Figure 2g,h). Notably, after two days of treatment, Exo-101 modulated the expression of population-specific transcription factors in CD4^+^ T-cells, with a trend towards the reduction of GATA-3, a significant decrease in T-bet and RORγt, and a significant increase in Foxp3 mRNA (Figure 2i-l). This effect was still visible after 6 days of treatment, for RORγt and Foxp3 expression.

**Figure 2:**
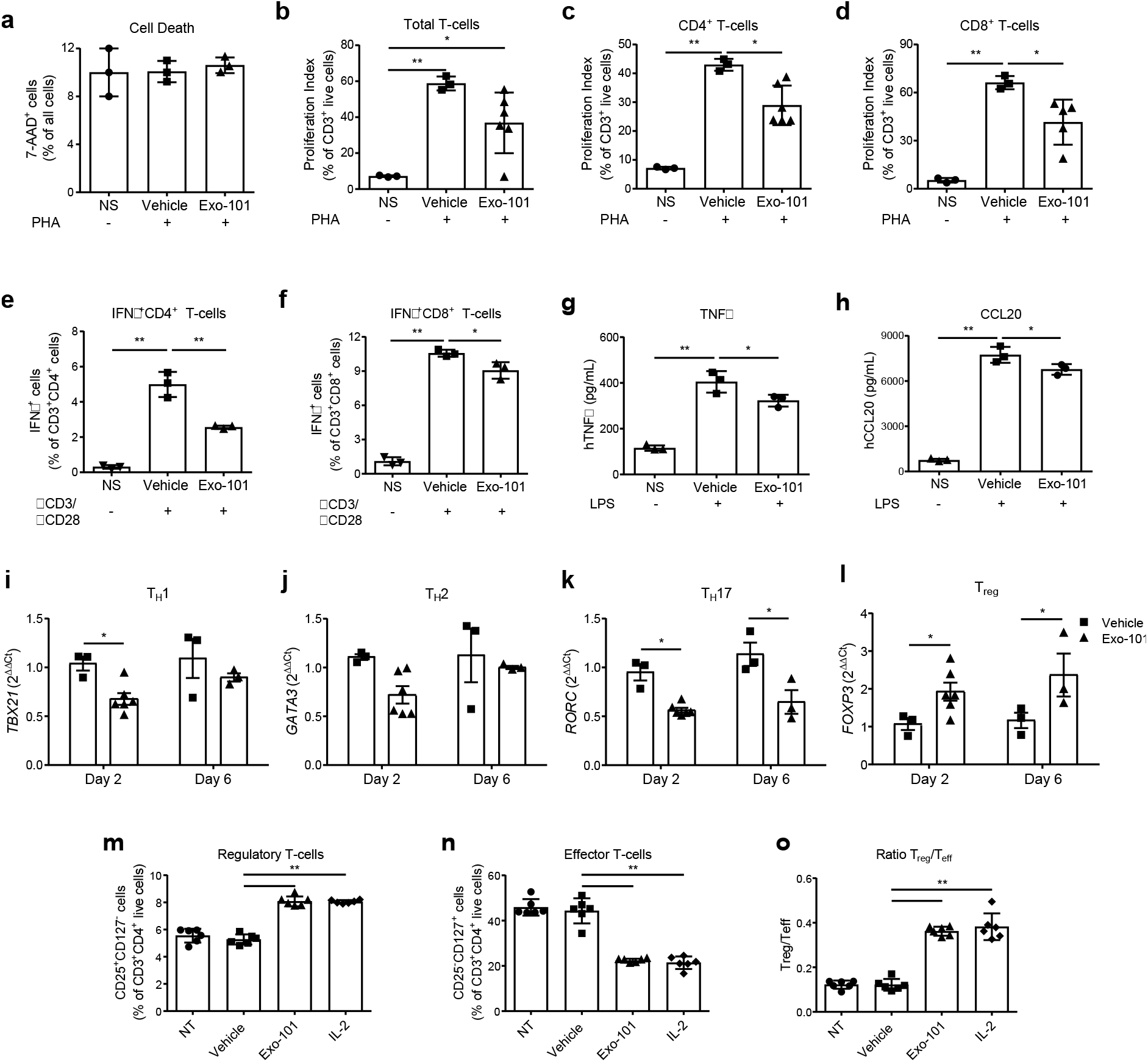
Exo-101’s immunomodulatory effects on T-cells *in vitro*. (**a-f**) Human PBMC, stimulated with PMA or αCD3/αCD28 as indicated, were incubated with a single dose of Exo-101 at 1×10^10^ particles/mL for 6 days, followed by flow cytometry analysis (n≥3). (**g, h**) TNFα and CCL20 release by LPS-stimulated PBMC after a 24h incubation with 1×10^10^ particles/mL Exo-101 (n=3). (**i-l**) Relative expression of transcription factors by PBMC incubated with Exo-101 for 2 or 6 days (n≥3). Cells were sorted based on FSC, SSC, CD3 and CD4, prior to RNA extraction. (**m-o**) Phenotyping of αCD3/αCD28-activated PBMC, after 6 days of incubation with Exo-101 at 1×10^10^ particles/mL or IL-2 (100 IU/ml) and TGF-β (5ng/mL) (n=6). All results are presented as mean +/- SD. *p≤0.05, **p≤0.01, ***p≤0.001, ****p≤0.0001. NS, non-stimulated; NT, non-treated.

To validate the differences in transcriptional regulation, we determined the populational T-cell changes by flow cytometry. As expected, Exo-101 treatment promoted the differentiation into regulatory T-cells (Treg), while inhibiting effector populations (Figure 2m-o). In this experiment, Tregs were defined as CD4^+^CD25^+^CD127^-^ cells, and the results were confirmed using a gating strategy based on the expression of the transcription factor Foxp3 (Figure S2). Importantly, Exo-101’s effect was shown to be as powerful as IL-2 for the induction of Treg (Figure 2m and S2).

### Exo-101 reduces the expression of psoriasis markers, in an *in vitro* 3D model

Considering the results from Figure 2, which evidenced an immunomodulatory effect of Exo-101, particularly focused on the Th17 and Treg response, we decided to evaluate the therapeutic benefit of Exo-101 in psoriasis. Using an in vitro 3D model of reconstructed human epidermis, engineered to be histologically and metabolically similar to psoriatic skin, we show that Exo-101 treatment significantly reduced the expression of the inflammatory mediators IL-6, IL-8, CXCL10 and COX-2, and had a tendential effect on *IFNG* and *TNFA* mRNA (Figure 3a-h). Moreover, the expression of psoriasin (S100A7) and beta-defensin-2 (DEFB4), two psoriasis-associated antimicrobial peptides, was also significantly decreased (Figure 3i,j).

**Figure 3:**
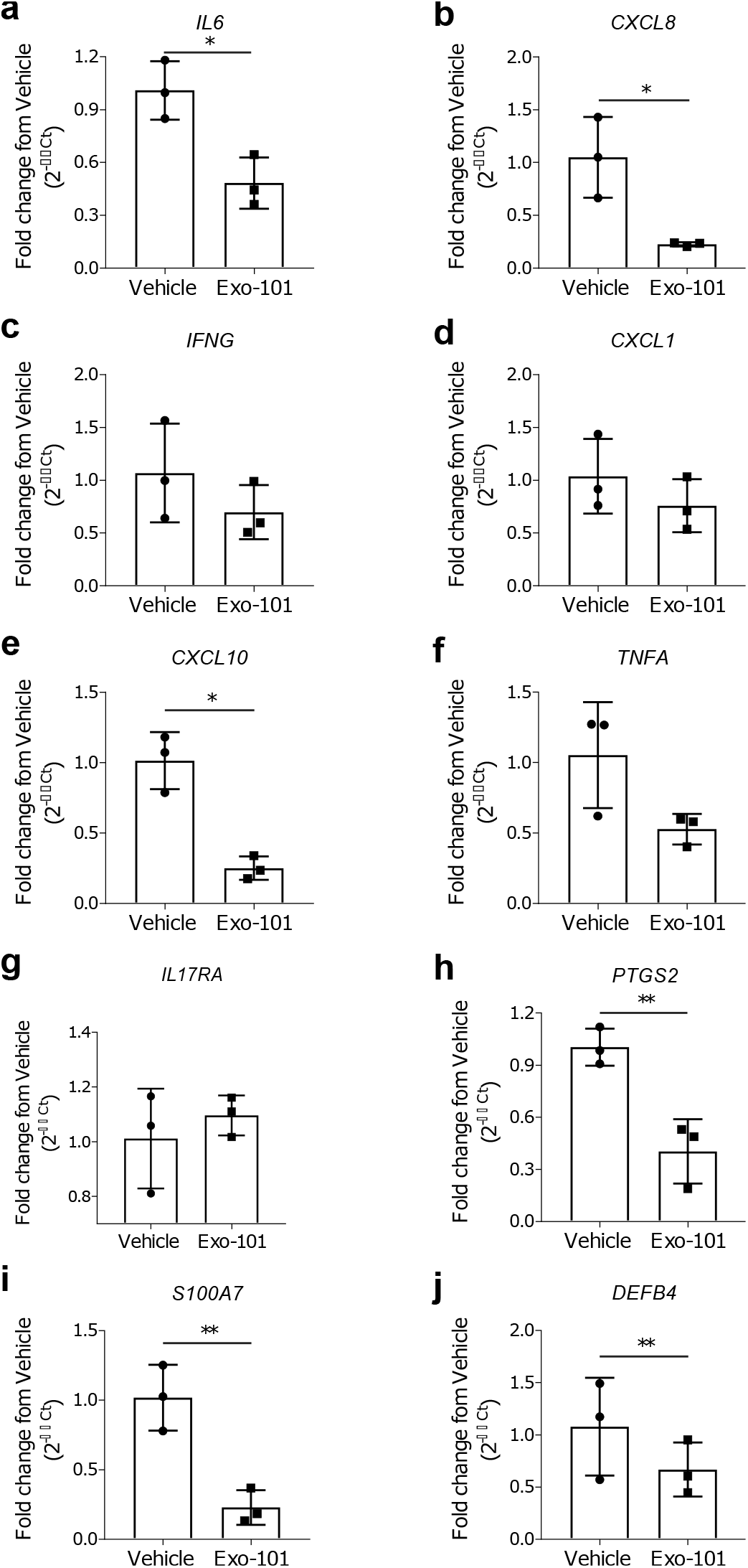
Decreased expression of epidermal psoriatic features after Exo-101 treatment. Relative expression of (**a-h**) pro-inflammatory mediators and (**i, j**) anti-microbial peptides by a 3D model of human psoriatic epidermis, treated with 1×10^10^ particles/mL Exo-101 twice daily for 6 days (n=3). All results are presented as mean +/- SD. *p≤0.05, **p≤0.01.

### Exo-101 shows a modest effect in imiquimod-induced psoriasis, regulating keratinocyte proliferation and T-cell homeostasis

Application of imiquimod, a TLR7 agonist, to mouse’s skin leads to the development of psoriatic features, such as, among others, epidermal thickening, erythema, inflammatory cell infiltration and epidermal expression of IL-17 (van der Fits et al. 2009). Daily topical treatment with Exo-101, dissolved in a slow-release hydrogel and applied one hour after imiquimod, did not significantly ameliorate macroscopic psoriasis-like features, when compared to hydrogel alone (Figure 4a,b). However, Exo-101 was significantly superior to hydrogel at reducing acanthosis, as seen by microphotographs (Figure 4c,d). The expression of the inflammatory markers TNFα, IFNγ, IL-17A, CCL20 and CXCL1, albeit significantly reduced when compared to the imiquimod control, was similar between the two groups receiving hydrogel, with or without Exo-101 (Figure 4e-i). When comparing the three treatment groups, there were no major changes in the skin infiltration of most inflammatory cells, including neutrophils, macrophages, total T-cells, γδ T-cells and total CD4^+^ T-cells (Figure 4j-k and S3). CD8^+^ T-cells were slightly reduced in Exo-101-treated animals compared to the skin of mice receiving only hydrogel (Figure 4l). Furthermore, Exo-101 tendentially increased the number of Treg in the skin of imiquimod-treated mice (Figure 4m), a finding that is consistent with gene expression data from diabetic mice with chronic skin wounds (Figure S4)(Cardoso et al. 2021; Henriques-Antunes et al. 2019). These findings indicate that, in a context of psoriasis, Exo-101 improves certain pathological features, possibly through a mechanism that involves both local and immune cells, but did not significantly reduce the overall disease burden in this model.

**Figure 4:**
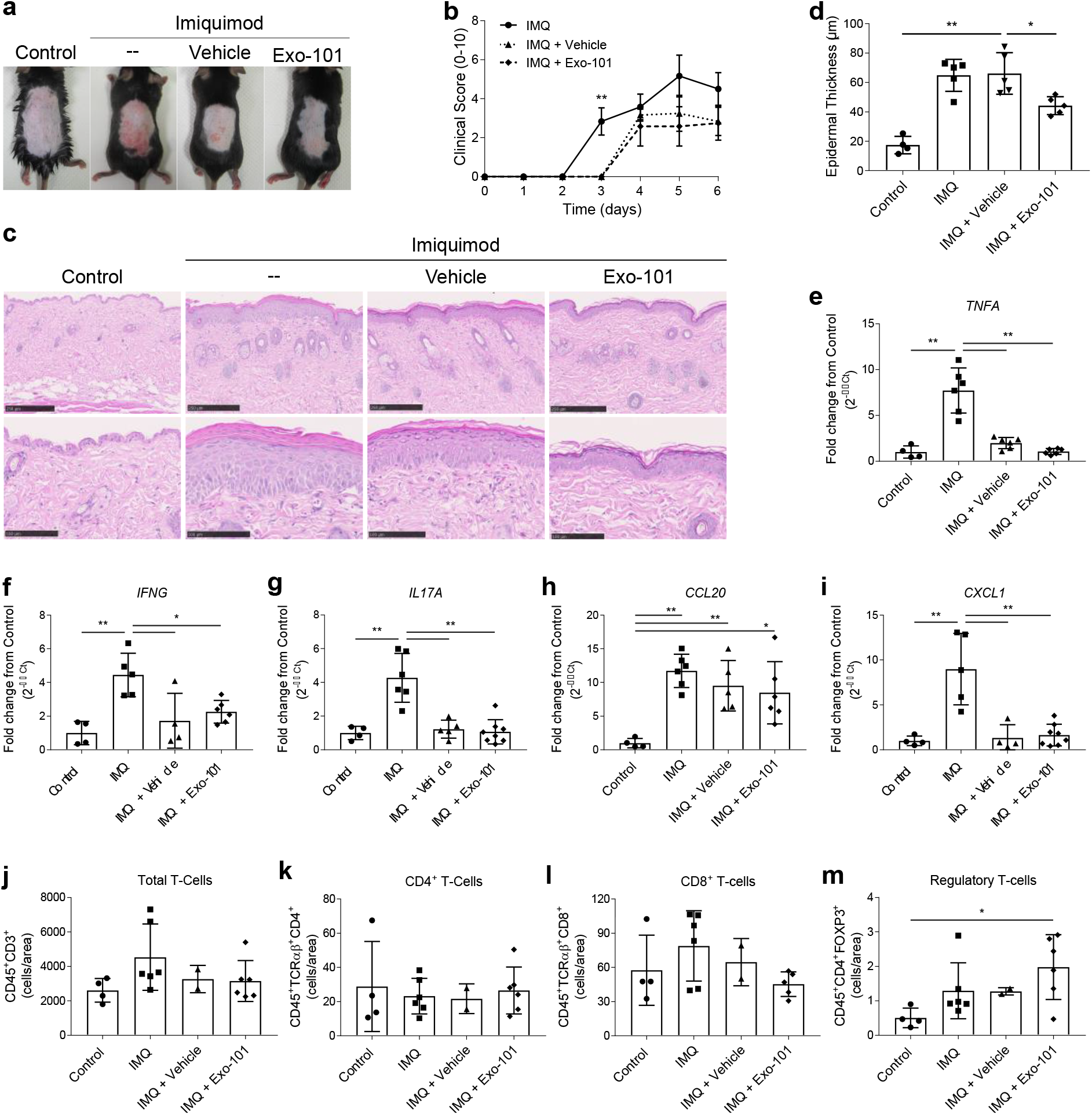
Therapeutic potential of Exo-101 for imiquimod-induced psoriasis. C57BL/6 mice received 5% imiquimod (IMQ) on their shaved backs (approx. 7 cm^2^), daily for 6 days. A topical formulation containing 2×10^10^ particles/mL Exo-101 was applied to the same area every day, 1h after IMQ. (**a**) Representative photos and (**b**) clinical score (n=6). (**c**) Representative H&E microphotographs used to measure (**d**) epidermal thickness (n≥4). Scale bars=250μm or 100μm, respectively for upper and lower microphotographs. Epidermal thickness was measured from *stratum basale* to *stratum granulosum*, averaging 5 measurements per section, for a total of 20 data points per animal. (**e-i**) Relative expression of pro-inflammatory mediators in the skin of all tested groups (n≥4). (**j-m**) Digested skin of all four test groups was analyzed by flow cytometry for identification of total, CD4^+^, CD8^+^ and regulatory T-cells (n≥2). All results are presented as mean +/- SD. *p≤0.05, **p≤0.01.

## DISCUSSION

Over the last years, sEV have been explored for their potential as cell-free immunomodulatory and regenerative agents. Indeed, unmodified or engineered sEV were shown to have therapeutic potential across multiple conditions, including cancer (Yuan et al. 2019), inflammatory lung diseases (Zhang et al. 2018; Zhu et al. 2014) and autoimmunity (Lai et al. 2019). Here, we explore the mechanism of action of sEV isolated from UCB-MNC, Exo-101, and evaluate their effect in psoriasis models.

*In vitro*, Exo-101 exhibits anti-inflammatory properties, affecting macrophage differentiation and cytokine production. These results are consistent with previous findings using sEV isolated from bone marrow (Cao et al. 2019) or cord blood (Hu et al. 2020). We also show that the presence of Exo-101-induced M2 macrophages has a down-stream effect on neighboring cells, such as skin fibroblasts, reducing their response to an inflammatory trigger. Moreover, Exo-101 strongly inhibited cell proliferation and cytokine production by LPS-stimulated total CD4^+^ and CD8^+^ T-cells, consistent with previous reports (Del Fattore et al. 2015; Di Trapani et al. 2016). This outcome is possibly due to an effect on T-cell differentiation, given that Exo-101 promotes a shift from a Th1 or Th17 into a Treg phenotype. Notably, Exo-101 stimulus was shown to be as effective as IL-2 in promoting Treg development.

Our *in vitro* findings evidenced a potential mechanism of action for Exo-101, responsible for a shift in the expression of transcription factors, which favor Treg differentiation and concomitantly silence Th17 signaling. Biologics targeting the Th17 axis (α-IL-17 and α-IL-23) have been proved to be clinically effective in ameliorating psoriasis symptoms (Gordon et al. 2018; Krueger et al. 2019; Papp et al. 2018). Additionally, previous reports suggest that Treg, a typically tolerogenic cell population, play a crucial role the maintenance of skin homeostasis. Treg-deficient animals present an exacerbated response to imiquimod (Hartwig et al. 2018) and Treg from psoriatic patients display an impaired suppressive function (Sugiyama et al. 2005). Hence, we hypothesized that Exo-101’s profile could be therapeutically beneficial in psoriasis. To test this, we employed a model of reconstructed human epidermis, composed of keratinocytes in various stages of differentiation, and pre-treated to display psoriasis-like inflammatory features. Exo-101 treatment significantly reduced the expression of psoriasis-associated molecules, including IL-6 and IL-8, as well as antimicrobial peptides S100A7 and DEFB4, thereby supporting its therapeutic potential for this disease.

In order to test Exo-101’s effect *in vivo*, we first designed a micelle-rich hydrogel that solidifies at normal body temperature, thus reducing product loss when applied to the skin. Psoriasis-like symptoms were induced by topical applications of imiquimod, and hydrogel was applied one hour later, alone or containing Exo-101 vesicles. In this *in vivo* model, Exo-101 proved superior to hydrogel in reducing or preventing keratinocyte hyperproliferation, as measured by epidermal thickness. Yet, there were no significant differences in the disease scores, mRNA expression and cellular profile between the two treatment groups. While it is possible that the application of hydrogel alone strongly improves psoriatic symptoms, the results observed are better explained by the possible trapping of imiquimod molecules in hydrogel micelles, therefore preventing full symptom development. An alternative experimental setting would either allow for a longer time interval between imiquimod and test treatment applications and/or require the increase of imiquimod dosage. Nevertheless, in line with previous *in vitro* data, Exo-101 had a positive effect on keratinocytes and was tendentially stronger than hydrogel alone in shifting skin cellular infiltrates toward a tolerogenic profile. Importantly, gene expression data from chronic wounds support the increase in Treg differentiation following Exo-101 treatment. Given the incomplete therapeutic response of imiquimod-treated animals to Exo-101, it is plausible that Exo-101 could act as an adjuvant treatment, in combination with standard therapies, such as anti-IL-17A or anti-IL23. This strategy would not only target immune-driven disease pathways, but also simultaneously stimulate repair mechanisms in the skin.

In conclusion, we show that Exo-101 decreases inflammation by targeting different cell populations, such keratinocytes, fibroblasts and macrophages, and by modulating T-cell differentiation and cytokine production. These findings warrant further proof-of-concept studies on the therapeutic potential of Exo-101 in inflammatory skin conditions, in particular, diseases thought to benefit from Th17/Treg-targeting.

## MATERIALS AND METHODS

### UCB collection, testing and data protection

Human UCB samples and relevant donor information were collected following signed informed consent, under approval of the Portuguese National Data Protection Committee and the ethics committees from five Portuguese hospitals, according to local legislation and following the principles of the Declaration of Helsinki. UCB processing, including microbiological testing, and storage were performed by an accredited biobank (Stemlab, S.A, Cantanhede, Portugal).

### Cell culture

#### UCB-MNC

Isolated UCB-MNC were cultured under 0.5% O_2_, for 18 hours, at a density of 2 million cells/mL in serum-free cell culture medium (Lonza AG, Basel, Switzerland) supplemented with 0.5 μg/mL FMS-like tyrosine kinase-3 (Peprotech, London, UK) and 0.5 μg/mL stem-cell factor (Peprotech, London, UK).

#### THP-1-derived macrophages

THP-1 cells (ATCC, Manassas, VA) were grown for 72 hours in the presence 25 nM PMA (Sigma, St. Louis, MO). THP-1-derived macrophages were then stimulated for 24 hours with 1 μg/mL LPS (Sigma, St. Louis, MO), followed by a 24-hour incubation with 1×10^10^ particles/mL Exo-101, when indicated.

#### Dermal fibroblasts

Normal human dermal fibroblasts were kept in Fibroblast Basal Medium, supplemented with Fibroblast Growth Kit Serum-Free, Phenol Red and Penicillin-Streptomycin-Amphotericin B Solution (ATCC, Manassas, VA). When appropriate, cells were counted after Hoescht staining (VWR International, Radnor, PA).

#### PBMC

Fresh human PBMC samples were isolated from volunteer donors, following informed consent, by density gradient centrifugation (Stemcell Technologies, Vancouver, Canada). For T-cell experiments, 2×10^5^ cells/well were activated with PMA (Sigma, St. Louis, MO), anti-CD3/-CD28 (see Table 2) or LPS (Sigma, St. Louis, MO), as indicated, followed by a single dose of Exo-101 at 1×10^10^ particles/mL.

### Exo-101 isolation

UCB-MNC culture media were subjected to an optimized isolation process, combining ultrafiltration and size exclusion chromatography (Cardoso et al. 2021). The resulting vesicles were characterized by transmission electron microscopy, flow cytometry, mass spectrometry for protein and lipid composition, RNA sequencing and nanoparticle tracking analysis, and found to be rich in CD63 and smaller than 200 nm (Cardoso et al. 2021).

### Gene expression

RNA was extracted with RNeasy Mini Kit (Qiagen, Hilden, Germany) and analyzed with total RNA chips in Bioanalyzer 2100 (Agilent Technologies, Santa Clara, CA). SuperScript™ IV VILO™ Master Mix (ThermoFisher Scientific, Waltham, MA) was used for reverse transcription and gene expression was detected by qPCR, using the primer pairs in Table 1.

**Table 1:**
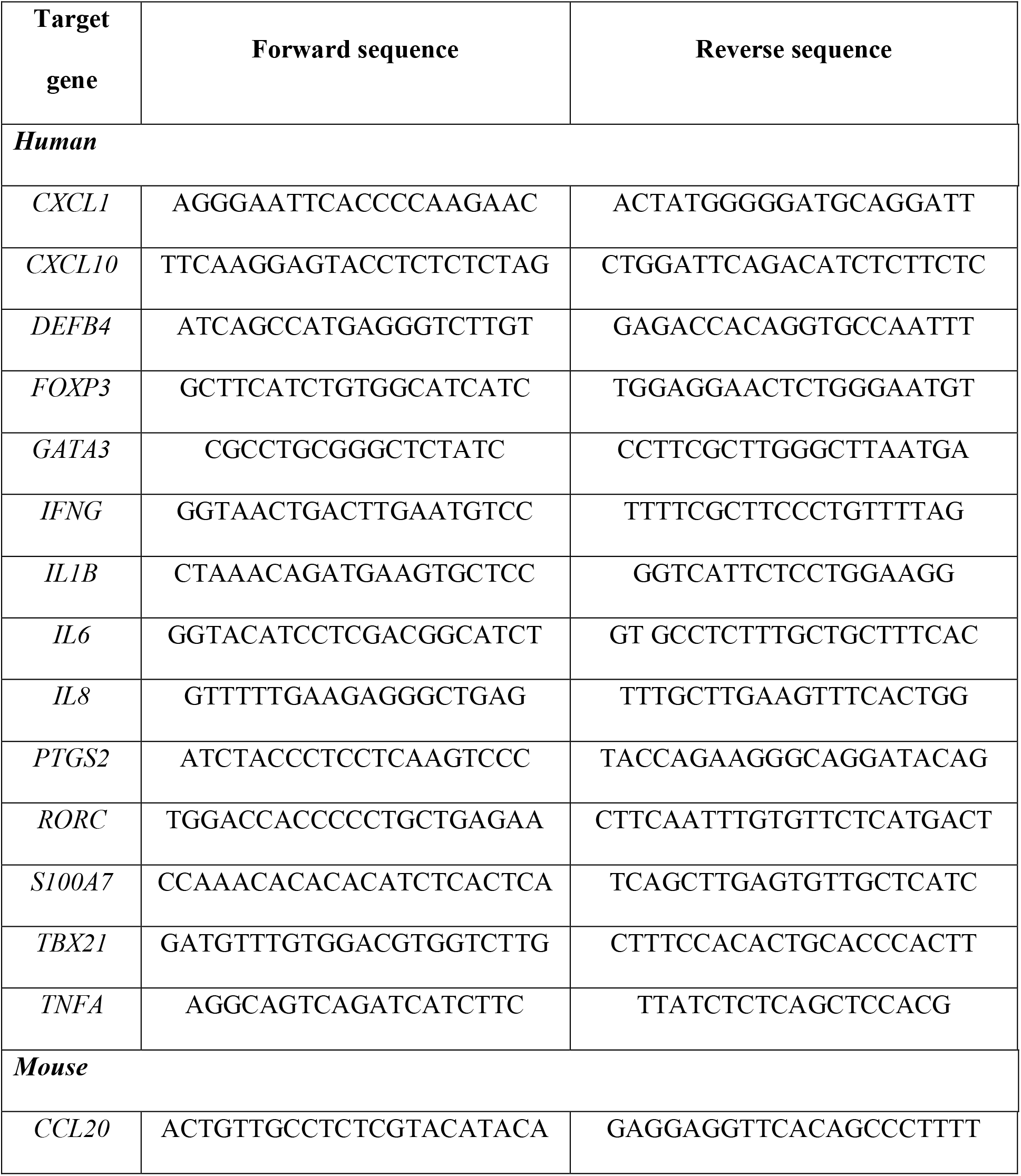
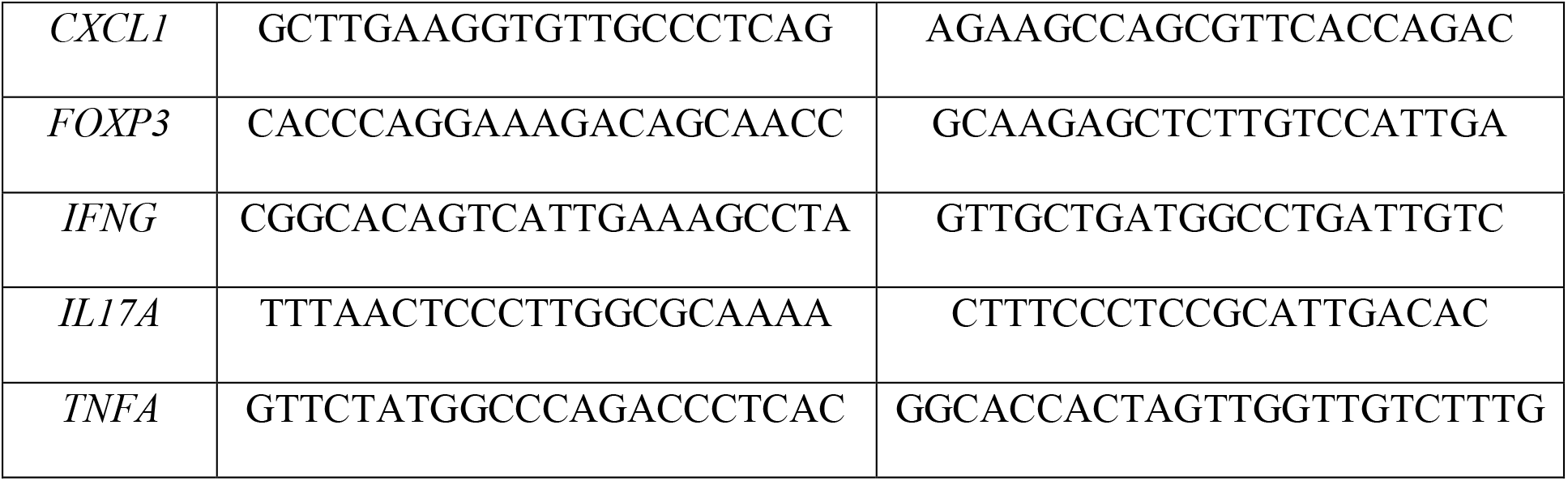
Human and mouse primer sequences for qPCR

**Table 2:** Human and mouse antibodies for flow cytometry

| Target | Clone | Manufacturer |
| --- | --- | --- |
| *Anti-human antibodies* |  |  |
| CD14 | M5E2 | BD Biosciences, San Jose, CA |
| CD163 | GHI/61 | BD Biosciences, San Jose, CA |
| CD28 | CD28.2 | BioLegend, San Diego, CA |
| CD3 | OKT3 | BioLegend, San Diego, CA |
| CD68 | Y1/82A | BD Biosciences, San Jose, CA |
| CD86 | FUN-1 | BD Biosciences, San Jose, CA |
| IL-2 | MQ1-17H12 | BioLegend, San Diego, CA |

| ***Anti-mouse antibodies*** |  |  |
| --- | --- | --- |
| CD11b | M1/70 | BD Biosciences, San Jose, CA |
| CD11c | HL3 | BD Biosciences, San Jose, CA |
| CD16/CD32 | 2.4G2 | Tonbo Biosciences, San Diego, CA |
| CD25 | PC61.5 | eBioscience, San Diego, CA |
| CD3e | 145-2C11 | BD Biosciences, San Jose, CA |
| CD4 | RM4-5 | BD Biosciences, San Jose, CA |
| CD45.2 | 104 | eBioscience, San Diego, CA |
| CD62L | MEL-14 | BD Biosciences, San Jose, CA |
| CD8 | 53-6.7 | BD Biosciences, San Jose, CA |
| Ly6C | AL-21 | BD Biosciences, San Jose, CA |
| Ly6G | 1A8 | BD Biosciences, San Jose, CA |
| TCR $\gamma/\delta$ | GL3 | BioLegend, San Diego, CA |
| TCR $\alpha/\beta$ | GL2 | BioLegend, San Diego, CA |

### Protein quantification and flow cytometry (human)

TNFα and CCL20 were measured by ELISA (BioLegend, San Diego, CA). M1 macrophages were defined as CD14^-^CD68^+^CD86^+^, and M2 macrophages as CD14^-^CD68^+^CD163^+^ by flow cytometry. Tregs were defined as CD4^+^CD25^+^CD127^-^ cells and the results were confirmed using a gating strategy based on the expression of the transcription factor *FOXP3* (Figure S2). Foxp3 expression was analyzed by flow cytometry using the Foxp3/Transcription Factor Staining Buffer Set (eBioscience, San Diego, CA). Whenever mentioned, IL-2 was used as a positive control for Treg development. Antibody details can be found in Table 2.

### Reconstructed psoriatic human epidermis

The 3D model of “psoriasis-like” human epidermis (Sterlab, Vallauris, France) was used according to the manufacturer’s instructions. Briefly, epidermal inserts were placed in a 12-well feeder-plate and allowed to equilibrate in the supplied culture medium for one day at 37°C and 5% CO_2_, before treatment with 1×10^10^ particles/mL Exo-101, twice a day for 5 consecutive days.

### Exo-101 formulation for *in vivo* topical use

Exo-101 is produced as a lyophilized powder. For *in vivo* experiments, Exo-101 was dissolved in a micellar hydrogel, which solidifies at body temperature, thereby reducing product loss when applied to the skin and allowing for a slow release of Exo-101 particles (not shown).

### Imiquimod-induced psoriasis

Animal experiments were approved by the ethical committee of the Spanish National Cardiovascular Research Center, and performed according to national and European regulations, respecting animal welfare guidelines and the 3R’s rule. 8-12-week-old C57BL/6 mice received daily applications of imiquimod (3M, Saint Paul, MN) on their shaved backs, for six consecutive days. One hour after every imiquimod application, 3×10^9^ particles/cm^2^ Exo-101 dissolved in hydrogel were delivered topically. Epidermal thickness was measured on H&E-stained samples, using the software NanoZoomer Digital Pathology NDP.view2 (Hamamatsu Photonics, Hamamatsu, Japan), from stratum basale to stratum granulosum, averaging 5 measurements per section, for a total of 20 data points per animal. For flow cytometry, skin was digested with 0.25 mg/mL Liberase (Roche, Basel, Switzerland) in serum-free RPMI, for 60 min at 37°C. Skin cell suspensions were stained with fluorescently-labelled antibodies, following Fc Block (Table 2). Absolute cell counts were performed using Trucount tubes (BD Biosciences, San Jose, CA). RNA analyses were performed as described above.

### Statistical analyses

Data analyses were performed with Prism 6 (GraphPad Software, San Diego, CA). Unpaired t-tests or one-way ANOVA were employed whenever appropriate (p<0.05).

## DATA AVAILABILITY

For any data or certificate requests, please contact Exogenus Therapeutics, S.A., at.

## ACKNOWLEDGMENTS

This work was co-funded by Regional Operational Program Center 2020, Portugal 2020 and European Union through FEDER within the scope of project CENTRO-01-0247-FEDER-022398. S.C.R.’s work was supported by FCT fellowship SFRH/BD/137633/2018. The authors greatly appreciated the assistance of Francisco Sanchez Madrid and Danay Cibrian for implementing the imiquimod-induced psoriasis mice model.

---

**Figure S1:**
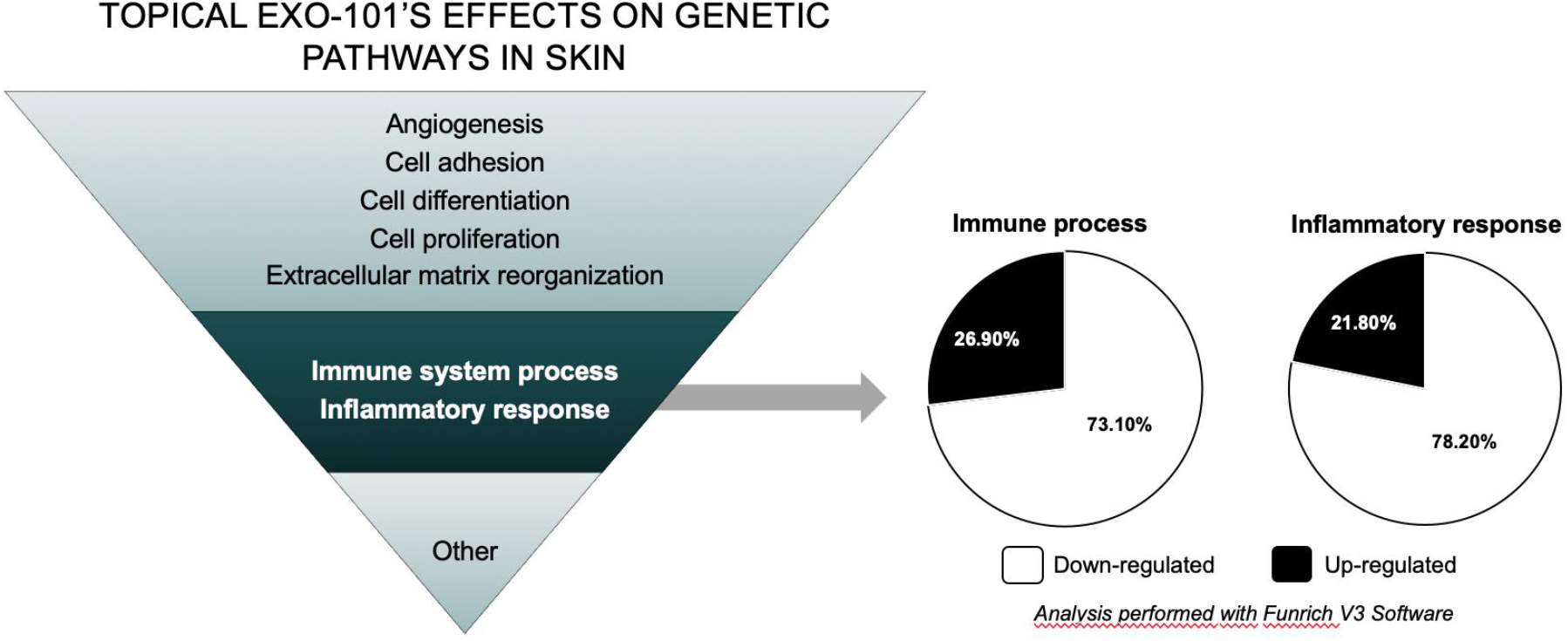
Effects of Exo-101 treatment on gene expression of chronic skin wounds. STZ-induced diabetic mice were treated topically with 2.5×10^9^ particles/mL Exo-101, twice per day for 15 days, after which skin mRNA was extracted. The inverted triangle on the left represents the different major processes affected by Exo-101 treatment. The circles on the right illustrate the percentage of inflammation- and immune system-related genes up- or down- regulated after Exo-101 treatment.

**Figure S2:**
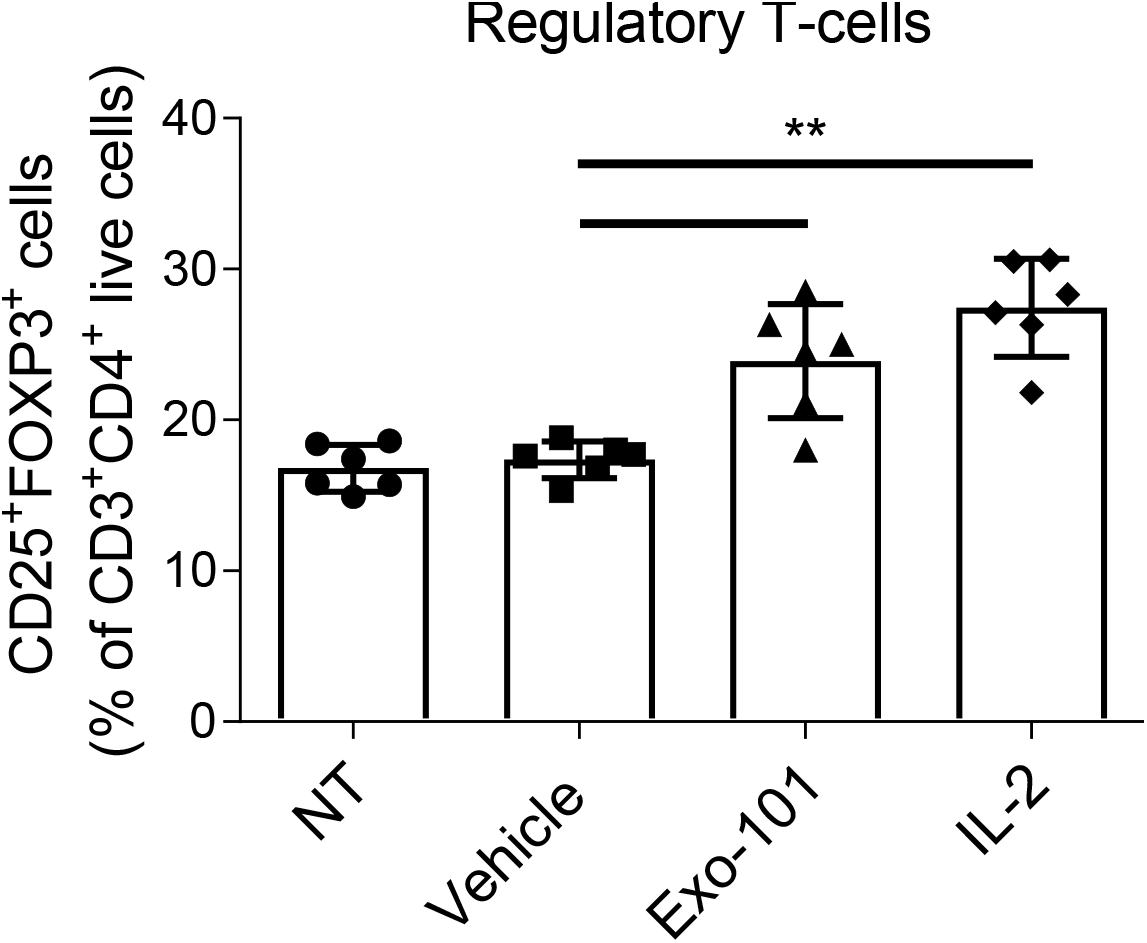
Effects of Exo-101 treatment on the percentage of regulatory T-cells (Treg). Human PBMC were activated with αCD3/αCD28 and treated for 6 days with Exo-101 (1×10^10^ particles/mL) or IL-2 (100 IU/ml) and TGF-β (5ng/mL) (n=6). Treg were defined as CD3^+^CD4^+^CD25^+^FOXP3^+^ cells. Comparison with Figure 1ab, where Treg were identified based on the expression markers CD25 and CD127. All results are presented as mean +/- SD. **p≤0.01. NT, non-treated.

**Figure S3:**
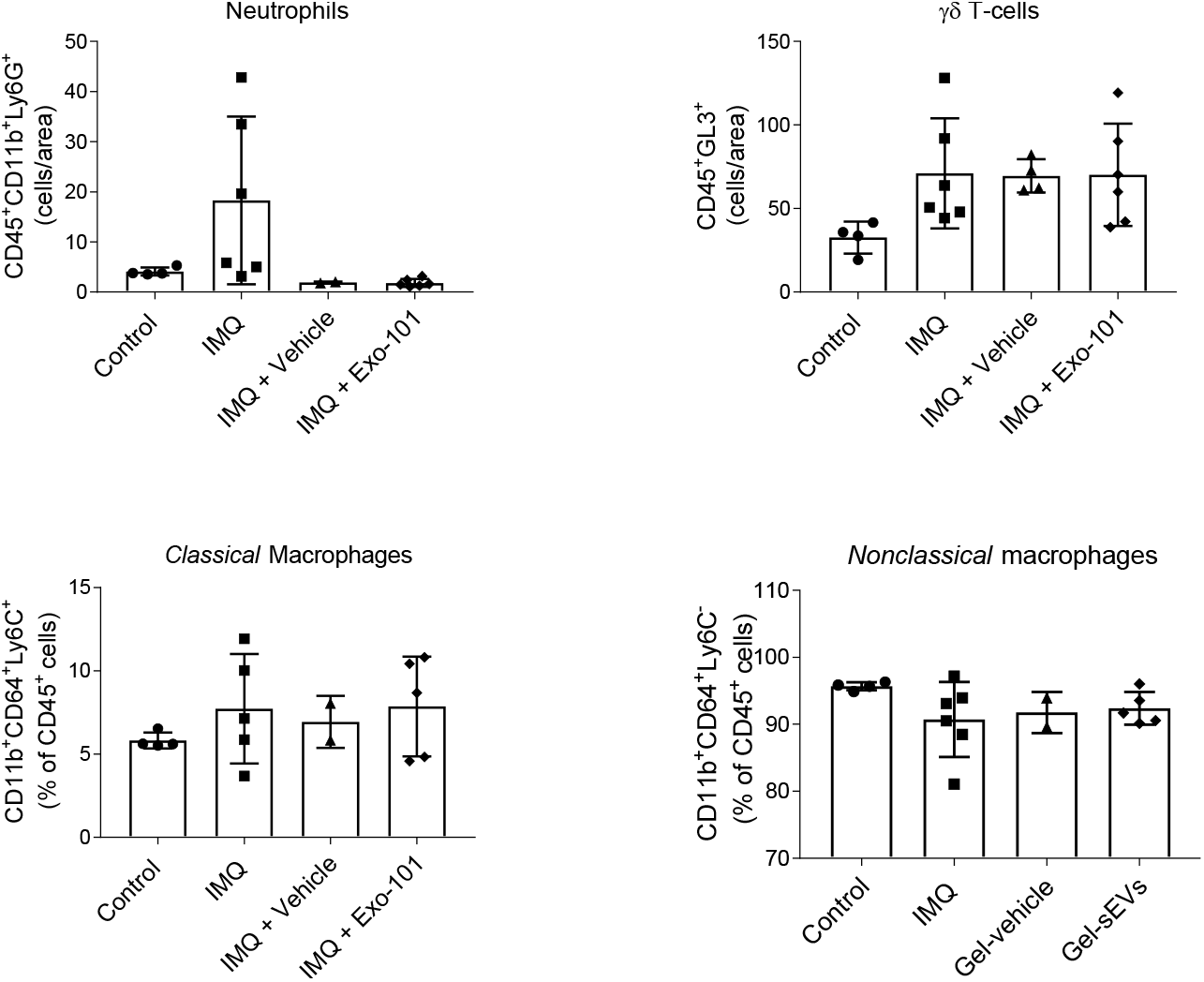
Cell populations in the skin of imiquimod- and Exo-101 treated animals. Flow cytometry identification of neutrophils, γδ T-cells and macrophages in digested skin of mice treated with imiquimod (IMQ) and/or Exo-101 as indicated (n≥2). All results are presented as mean +/- SD.

**Figure S4:**
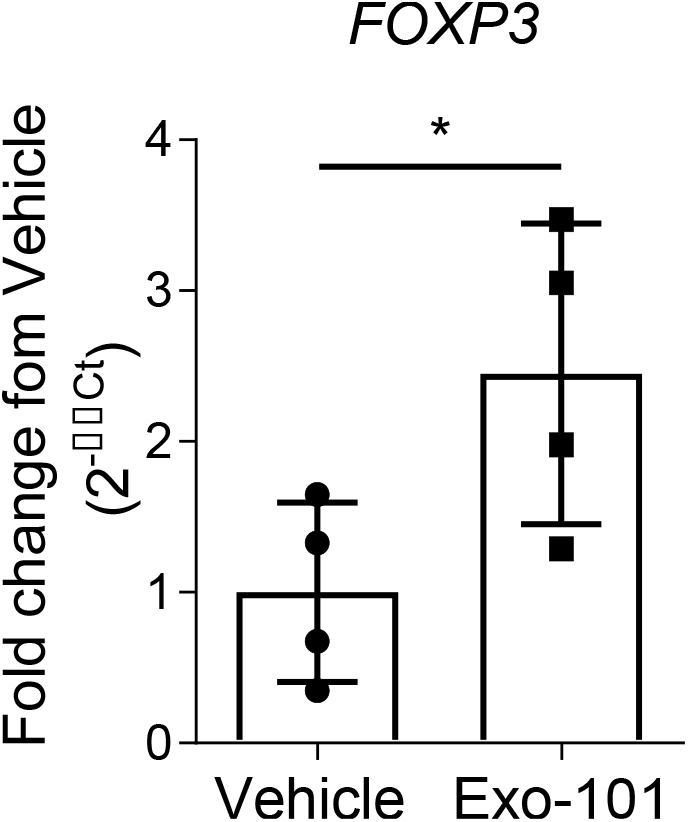
FOXP3 expression in the skin of Exo-101 treated mice with chronic wounds. STZ-induced diabetic mice (n=4) were treated topically with 2.5×10^9^ particles/mL Exo-101 or vehicle, twice per day for 15 days, after which skin mRNA was extracted. All results are presented as mean +/- SD. *p≤0.05.

